# Human Genes Are *In Silico* Potential Targets For Rice miRNA

**DOI:** 10.1101/2020.01.02.893040

**Authors:** Aizhan Rakhmetullina, Anna Pyrkova, Dana Aisina, Anatoliy Ivashchenko

**Affiliations:** Department of Biotechnology, SRI of Biology and Biotechnology Problems, al-Farabi Kazakh National University, Almaty, al-Farabi 71, Almaty, 050040, Kazakhstan

## Abstract

Exogenous miRNAs enter the human body through food, and their effects on metabolic processes can be considerable. It is important to determine which miRNAs from plants affect the expression of human genes and the extent of their influence. The binding sites of 738 osa-miRNAs that interact with 17508 mRNAs of human genes were determined using the MirTarget program. The characteristics of the binding of 46 single osa-miRNAs to 86 mRNAs of human genes with a value of free energy (ΔG) interaction equal 94% to 100% from maximum ΔG were established. The findings showed that osa-miR2102-5p, osa-miR5075-3p, osa-miR2097-5p, osa-miR2919 targeted the largest number of genes at 38, 36, 23, 19 sites, respectively. mRNAs of 86 human genes were identified as targets for 93 osa-miRNAs of all family osa-miRNAs with ΔG values equal 94% to 98% from maximum ΔG. Each miRNA of the osa-miR156-5p, osa-miR164-5p, osa-miR168-5p, osa-miR395-3p, osa-miR396-3p, osa-miR396-5p, osa-miR444-3p, osa-miR529-3p, osa-miR1846-3p, osa-miR2907-3p families had binding sites in mRNAs of several human target genes. The binding sites of osa-miRNAs in mRNAs of the target genes for each family of osa-miRNAs were conserved when compared to flanking nucleotide sequences. mRNA human genes of osa-miRNAs are candidate genes of cancer, cardiovascular and neurodegenerative diseases.

## Introduction

Plant miRNA (pl-miR) and animal miRNA (an-miR) are exogenous miRNA or xeno miRNA (ex-miR or xe-miR) for humans and do not have distinctive features among themselves. Therefore, in the human body, they will be perceived as the general diversity of the endogenous miRNA (en-miR) of a human, and all human genes can potentially be their targets. The effect of ex-miR on human target genes and the consequences of this effect will depend on their concentration and the duration of their presence in the cells. pl-miR can enter the human body through the gastrointestinal tract and spread with blood in combination with proteins or as part of exosomes (Buck et al., 2014; Escrevente et al., 2011; Montecalvo et al., 2012; van der Grein & Nolte-’t Hoen, 2014). Many studies have shown that miRNAs that enter the gastrointestinal tract with food (dietary miRNA) are then found in various tissues (Chiang et al., 2015; Jonathan et al., 2013; Liang et al., 2015; Stephen & Snow, 2017; Vaucheret & Chupeau, 2012; Zempleni et al., 2017; Zhang et al., 2019; Zhang et al., 2012; Zhao et al., 2018). For example, аfter consumption of a fresh maize diet most zma-miRNAs were detected in the heart, brain, mammary gland, lung, liver, kidney and serum exosomes of pig (Luo et al., 2017). The concentration of pl-miR in different human organs varies and can be compared with en-miR. Some studies have found minor concentrations of pl-miR in humans and animals (Zhang et al., 2016). The inclusion of pl-miR into the recipient’s organism causes reproducible changes of some properties and physiological processes in it (Cui et al., 2017; Javed et al., 2017; Lang et al., 2019; Vaucheret et al., 2012; Zhang et al., 2012). ex-miR involvement in the regulation of recipient gene expression may affect disease (Chin, Fong & Somlo, 2016; Gopinath, 2019; Hou et al., 2018; Jones et al., 2016). The program for searching binding sites in target genes can reliably identify pl-miR and an-miR binding sites regardless of the origin of miRNAs (Ivashchenko et al., 2016), which makes it possible to predict the interaction of pl-miR with hsa-mRNA. We studied the possible interactions of pl-miRs with hsa-mRNA genes after pl-miRs have entered the human body. It was also assumed that pl-miRs can circulate in the blood throughout the body in the absence of features restricting their entry into any cell (Wagner et al., 2015). The basis for this assumption is the diversity of the nucleotide composition of hsa-mRNAs, some of which overlap with diverse pl-miRs (Liang et al., 2013; Pirro et al., 2016).

Recently, the miRNAs of plants ingested for food ex-miR have been actively studied in the regulation of vital processes in humans and animals (Arroyo et al., 2011; Liang et al., 2012; Luo et al., 2017; Vaucheret & Chupeau, 2012; Zhang et al., 2016). It is presumed that ex-miRs regulate cellular function in healthy cells and act as important mediators in the development of animal diseases (Cong et al., 2018; Hoy & Buck, 2012; Makarova et al., 2016; Rutter & Innes, 2018). Ex-miRs can function as evolutionary linkers between different species and contribute to signal transmission both within and between species (Arteaga-Vazquez et al., 2006; Axtell, Westholm & Lai, 2011; Millar & Waterhouse, 2005; Moran et al., 2017; Zhao, Cong & Lukiw, 2018). Based on the available data, the authors suggest that such xe-miRNAs contribute to the beneficial properties of medicinal plants (Lukasik & Zielenkiewicz, 2016; Xie, Weng & Melzig, 2016), contribute to the negative properties of disease-causing or poisonous plants, and cross-link species between kingdoms of living organisms by participating in many of the mechanisms associated with the occurrence and pathogenesis of various diseases (Malloci et al., 2018; Melnik, John & Schmitz, 2014; Perge et al., 2017; Pogue et al., 2014).

To determine which pl-miRs affect the mRNAs of human genes, we chose osa-miRNAs because rice has the most miRNAs in the plants, and rice is the most common source of human nutrition. Most plants contain well-known miRNAs, which serve as typical regulators of plant growth and development (Bari, Orazova & Ivashchenko, 2013; Bari et al., 2014, Nair et al., 2010).

## Results

### Characteristics of the interaction of single osa-miRNAs with mRNA of human genes

Currently, 738 miRNAs encoded by the rice genome are known. For these osa-miRNAs, target genes from among 17508 human genes were searched. A total of 82 miRNAs with one to four target genes ware identified (Table S1). miR11339-3p and miR11339-5p; miR1425-3p and miR1425-5p; miR1432-3p and miR1432-5p; miR1870-3p and miR1870-5p; miR2096-3p and miR2096-5p; miR2867-3p and miR2867-5p; and miR390-3p and miR390-5p, originating from the same pre-miRNA, had binding sites in the mRNAs of different genes. The functions of the 162 identified target genes were diverse.

In the group of 49 miRNAs with five or more target genes, there were several miR-3p/miR-5p pairs that originated from the same pre-miRNA (Table S2). The total number of target genes for miRNAs with five or more genes was 479. The number of target genes for miR408-3p, miR5150-3p, miR528-3p, and miR530-3p was comparable to the number of target genes for miR408-5p, miR5150-5p, miR528-5p, and miR530-5p. For miR1847.1-5p, miR1850.1-5p, miR2094-5p, miR2097-5p, miR2102-5p, and miR3979-5p, the set of target genes was significantly larger than that for each corresponding miRNA-3p. Only miR5144-3p had four-fold more target genes compared to the number attributed to miR5144-5p. The miRNAs with the largest number of target genes were miR2102-5p (38 genes), miR5075-3p (36 genes), miR2097-5p (23 genes), and miR2919 (19 genes). Consequently, at high concentrations, these miRNAs could significantly change the metabolism of recipient human cells.

A total of 641 target genes were identified for 131 single miRNAs, which is approximately 3.7% of the total number of studied human genes.

Table 1 shows the characteristics of the binding of some osa-miRNAs with mRNAs of human genes. Each of the 35 miRNAs could bind to mRNAs of one target gene, six miRNAs had targets with two genes, and four miRNAs had three target genes with a value ΔG/ΔGm equal to 94-98%. miR2102-5p had 11 target genes with a value ΔG/ΔGm of 94-100%, and the free energy of the interaction of the miRNAs with the mRNAs of these genes varied from −115 kJ/mole to −121 kJ/mole. The miR2102-5p binding sites were located mainly in the 5’UTR, which suggests that they have a role in the early inhibition of the translation process. However, this property of miR2102-5p indicates the need to control its plant food-derived concentration in the human body. 19 target genes were associated with miR2919. miR5075-3p could bind to three mRNAs at binding sites located in the coding domain sequence and the 5’-untranslated region. The high-affinity binding sites were located in the 5’UTR and CDS of the mRNAs with the ΔG/ΔGm value was 94-98%. 17 miRNA binding sites were located in 5’UTR, 39 in CDS and 33 in 3’-untranslated region. Therefore, monitoring the concentrations of miR2102-5p, miR2919 and miR5075-3p in human biological fluids is also necessary.

**Table 1.**
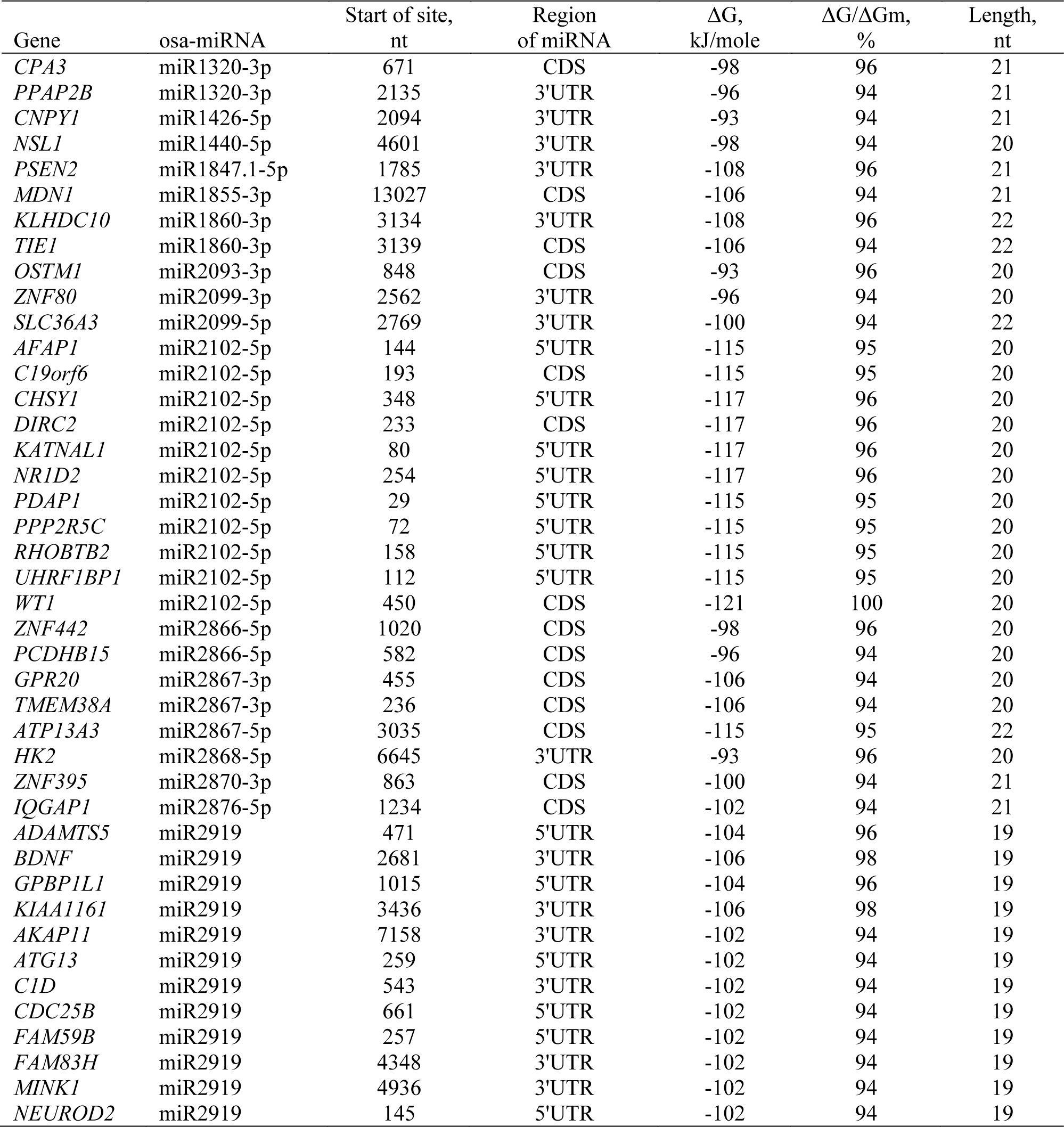
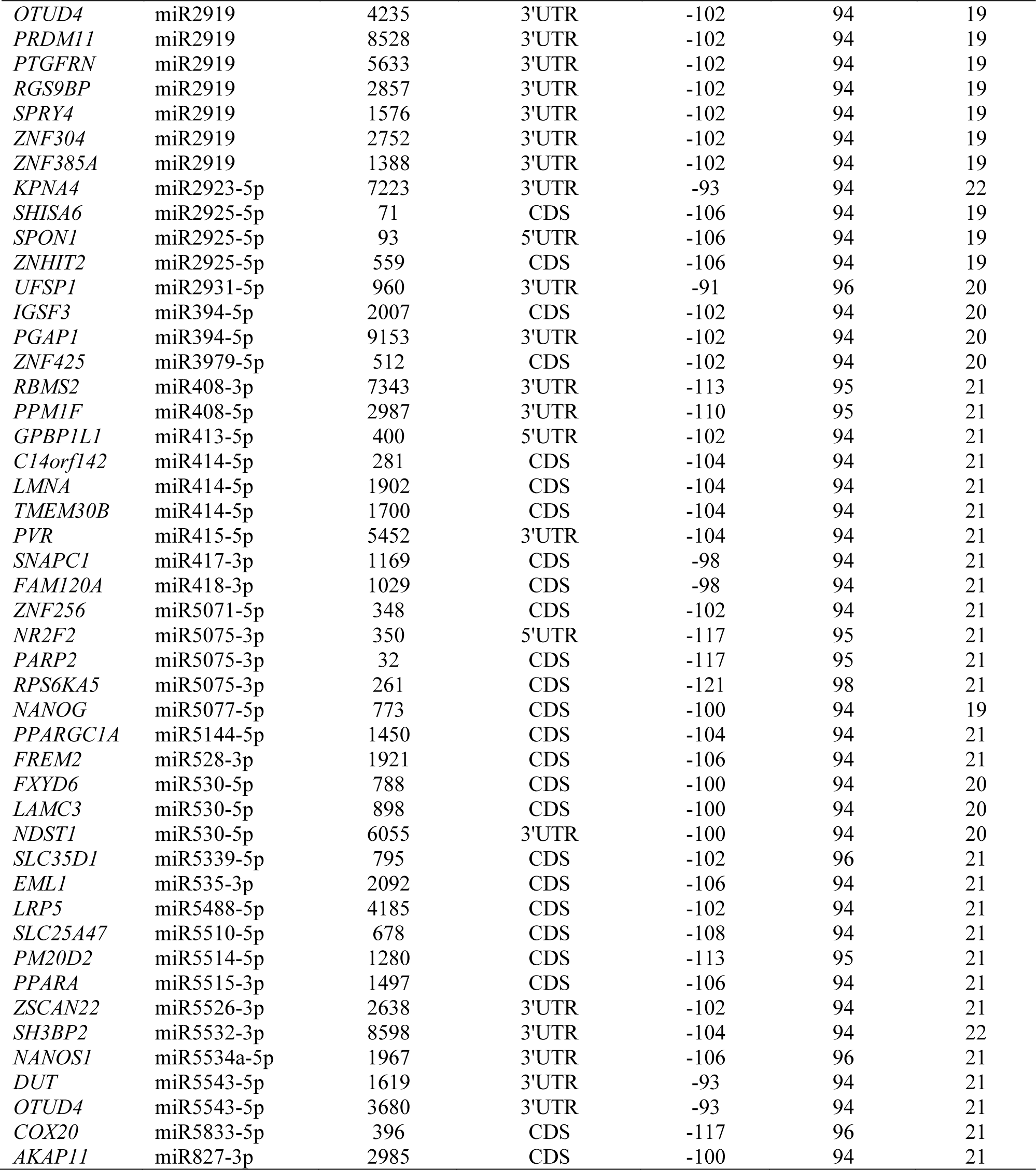
Characteristics of Interaction single osa-miRNAs with mRNA of Human Genes

Table S1-S3 show information about the genes targeted by plant miRNAs that may be involved in the development of various diseases. Most target genes are involved in the development of cancer of various types: *ENAH, MAPT, PRKCE, PRRT2, RBMS2, RHOBTB2, RPS6KA5 and ZFHX3*-breast cancer (Aisina et al., 2019); *ADAMTS5, PRAPRGG1A, PVR, SPRY4*-colorectal cancer; *CHSY1*-colorectal cancer and hepatocellular carcinoma; *PPM1F, FXYD6, DUT, FAM83H*-hepatocellular carcinoma; *HK2*-gallbladder cancer and leukaemia; *C19orf6*-ovarian carcinoma; *AFAP1*-oesophageal adenocarcinoma; *IOGAP1*-pancreatic ductal adenocarcinoma; *PPP2R5C*-lung adenocarcinoma; *PDAP1*-leukemia; *AKAP11*, *OSTM1, LRP5*-osteopetrosis; *WT1*-ovarian cancer and myeloid leukaemia; *NR1D2*-various cancers, including glioblastoma; CDC25B, *DIRC2*-renal carcinoma; and *UHRF1BP1*-cell carcinoma of the head and neck. Some genes are associated with other diseases: *NR2F2*-metabolic gene regulation and congenital heart defect; *PRAPA*, *UFSP1*-with increased cardiovascular disease; and *ATP13A3*-pulmonary tumour arterial hypertension and psychiatric disorder. *BDNF* has an important role in the neurogenesis and neuroplasticity of the brain; *PSEN2* and *LMNA* are associated with Alzheimer’s disease; *ZNF442* has a role in psychiatric disorders; and *NANOS1* is associated with retinoblastoma tumours. The list of oncological diseases caused by the target genes of the osa-miRNAs indicates that miRNAs can participate in the development of cancer not only in the gastrointestinal tract but also in other organs. Therefore, miRNAs that are ingested with food can be transferred to other tissues and organs.

The interaction of miRNAs and mRNA nucleotides of target genes shows how effectively these molecules bind. The schemes presented in Fig. 1 show the formation of hydrogen bonds between all the nucleotides of miR5075-3p, miR2866-5p, and miR2919 and the binding sites in mRNA. Because the MirTarget program takes into account the interaction of the noncanonical pairs A–C and G–U, it can be seen that the interaction of miRNAs and mRNAs preserves the spiral structures of both molecules, and therefore, stacking interactions are found between all the nucleotides of the miRNA and mRNA, which stabilize the duplex. miRNA binding sites are located in the 5’UTR, CDS, and 3’UTR.

**Figure 1.**
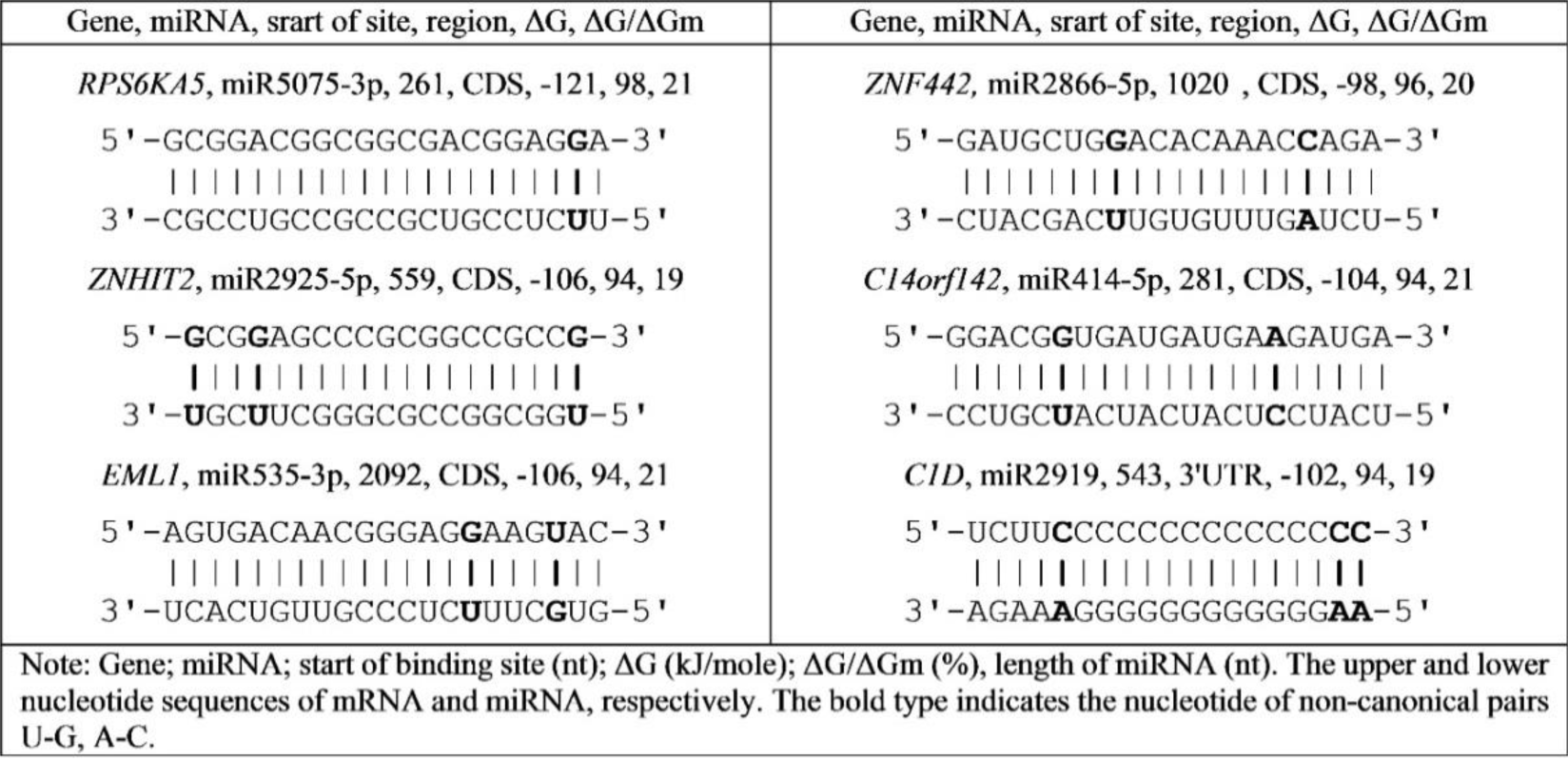
Schemes of the interaction of the nucleotide sequences of osa-miRNA with mRNA human genes.

One form of evidence for the reliability of miRNA interaction with mRNA is the establishment of a conservative nucleotide sequence of binding sites in mRNA of target genes (Atambayeva et al., 2017; Bari, Orazova & Ivashchenko, 2013; Bari et al., 2014; Yurikova et al., 2019). The results of the analysis of the similarity of the nucleotide sequences of the binding sites in mRNA of the target genes miR2102-5p, miR2919 and miR5075-3p are shown in Fig. 2. For all miR2102-5p, miR2919, and miR5075-3p binding sites, nucleotide conservation is present compared to the flanking nucleotides of mRNA target genes. With the complete complementarity of the nucleotides miRNA and mRNA (ΔG/ΔGm = 100%) of the target genes, absolute conservatism of the site along the entire binding site should be observed, as previously shown (Atambayeva et al., 2017; Bari, Orazova & Ivashchenko, 2013; Bari et al., 2014; Yurikova et al., 2019). When the ΔG/ΔGm value changes from 94% to 100% (Table 1), the basis for the interaction of the nucleotides miR2102-5p, miR2919 and miR5075-3p and mRNA are G-С pairs.

**Figure 2.**
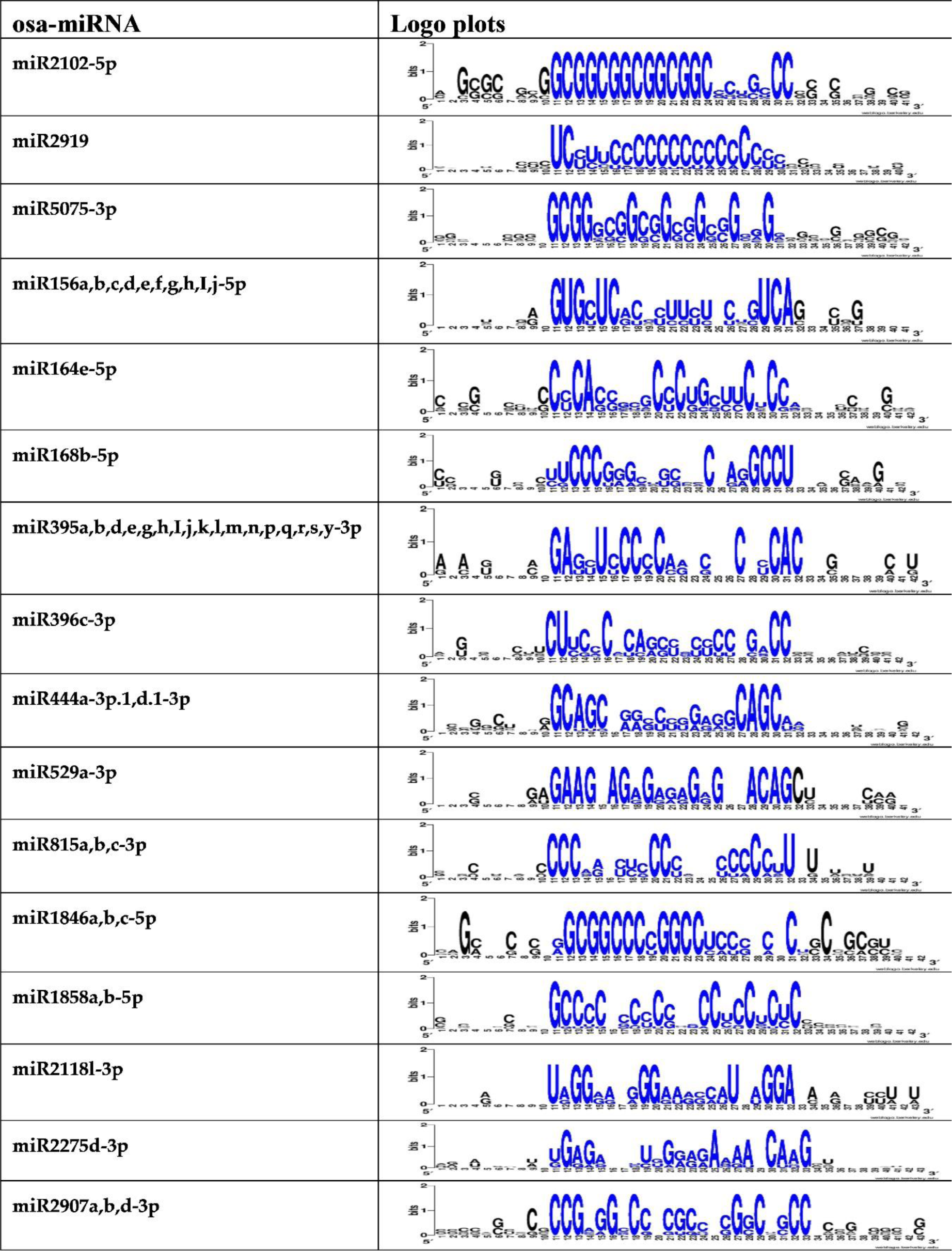
Logo plots of the nucleotide sequence variability of mRNA regions of human genes containing the binding sites of osa-miRNA.

### Characteristics of the interaction of osa-miRNA families with mRNAs of human genes

The total number of osa-miRNAs of all families is 146, and the number of their target genes is equal to 301, which is 1.7% of 17,508 studied human genes. The characteristics of the interaction of 93 osa-miRNAs of all family osa-miRNAs to 86 mRNAs of human genes with values from 94% to 98% were established (Table 2). miRNA of families such as miR156b-3p, miR159a.1,b,f-3p, miR164a,b,c,d,f-5p, miR166a,e-5p, miR166b,c,d,h-5p, miR167a,b,c,d,e,f,g,h,i,j-5p, miR172a,d-3p miR172c-3p, miR396d, miR396a,b-3p, miR531b-5p, miR815a,b,c-3p, miR1428b,c,d,e-3p and miR1858a,b-5p, has one target gene per family. All members of each miRNA family bind at one site due to the homology of their nucleotide sequences. Therefore, the expression of the target gene for each miRNA family will depend on the total concentration of all miRNA families. The miRNA families miR167e,i-3p, miR168b-5p, miR1846a,b,c-5p and miR2907a,b,c,d-3p had two target genes. miR2907a,b,c,d-3p has target genes *IRAK2* and *SLC25A37* with mRNAs of which these miRNAs interact with a large free energy of −123 kJ/mole and −125 kJ/mole, respectively. miR444b.1,c.1-3p had three target genes (*C19orf57, KAZN, NRG1*).

**Table 2.** Characteristics of interactions of osa-miRNA of families in mRNA of human genes

| Gene | osa-miRNA | Start of site, nt | Region of miRNA | $\Delta G$, kJ/mole | $\Delta G/\Delta G_m$, % | Length, nt |
| --- | --- | --- | --- | --- | --- | --- |
| *ANKRD27* | miR1428b,c,d,e-3p | 244 | CDS | -98 | 96 | 21 |
| *AP2A2* | miR156a,b,c,d,e,f,g,h,i,j,k-5p | 3797 | 3'UTR | -104 | 94 | 21 |
| *OBFC1* | miR156a,b,c,d,e,f,g,h,i,j-5p | 5427 | 3'UTR | -100 | 94 | 20 |
| *ZNF652* | miR156a,b,c,d,e,f,g,h,i,j-5p | 8125 | 3'UTR | -100 | 94 | 20 |
| *MROH2B* | miR156b-3p | 4615 | CDS | -106 | 94 | 21 |
| *PKHD1* | miR159a.1,b,f-3p | 9966 | CDS | -108 | 98 | 21 |
| *ASXL1* | miR164a,b,c,d,f-5p | 1923 | CDS | -110 | 95 | 21 |
| *CHST11* | miR166a,e-5p | 986 | CDS | -104 | 94 | 21 |
| *CCNY* | miR166b,c,d,h-5p | 367 | CDS | -110 | 95 | 21 |
| *PRICKLE2* | miR167a,b,c,d,e,f,g,h,i,j-5p | 1959 | CDS | -106 | 94 | 21 |
| *IL17RB* | miR167e,i-3p | 137 | CDS | -102 | 94 | 21 |
| *KIAA0528* | miR167e,i-3p | 523 | CDS | -102 | 94 | 21 |
| *APOBR* | miR168b-5p | 55 | CDS | -110 | 95 | 21 |
| *KLF14* | miR168b-5p | 623 | CDS | -115 | 98 | 21 |
| *GPR31* | miR172a,d-3p | 590 | CDS | -100 | 94 | 21 |
| *ATP12A* | miR172c-3p | 3557 | 3'UTR | -102 | 94 | 21 |
| *EIF3B* | miR1846a,b,c-5p | 2194 | CDS | -119 | 97 | 21 |
| *FASN* | miR1846a,b,c-5p | 11 | 5'UTR | -117 | 95 | 21 |
| *C20orf27* | miR1858a,b-5p | 921 | 3'UTR | -115 | 95 | 21 |
| *IRAK2* | miR2907a,b,d-3p | 1488 | CDS | -123 | 94 | 22 |
| *SLC25A37* | miR2907a,b,d-3p | 884 | CDS | -125 | 95 | 22 |
| *EDN1* | miR395a-q,t,y-3p | 1136 | 3'UTR | -108 | 96 | 21 |
| *CD22* | miR395b,d,e,g,h-t,y-3p | 2684 | 3'UTR | -104 | 94 | 21 |
| *CMKLR1* | miR396a,b-3p | 3970 | 3'UTR | -98 | 96 | 20 |
| *SCAMP5* | miR396a,b-5p | 1727 | 3'UTR | -102 | 94 | 21 |
| *SYVN1* | miR396a,b-5p | 854 | CDS | -102 | 94 | 21 |
| *AKAP13* | miR396d | 1736 | CDS | -100 | 94 | 20 |
| *RHBDF1* | miR444a-3p.1, d.1-3p | 1071 | CDS | -108 | 94 | 21 |
| *RUNDC1* | miR444a-3p.1, d.1-3p | 1191 | CDS | -108 | 94 | 21 |
| *CEP250* | miR444a-3p.2, b.2, c.2, d.2,e-3p | 1939 | CDS | -102 | 94 | 21 |
| *C19orf57* | miR444b.1, c.1-3p | 714 | CDS | -106 | 94 | 21 |
| *KAZN* | miR444b.1, c.1-3p | 4874 | 3'UTR | -106 | 94 | 21 |
| *NRG1* | miR444b.1,c.1-3p | 2444 | 3'UTR | -106 | 94 | 21 |
| *ATP6V0A4* | miR529a-3p | 241 | 5'UTR | -100 | 94 | 20 |
| *CCDC94* | miR529a-3p | 379 | CDS | -100 | 94 | 20 |
| *CHD2* | miR529a-3p | 6684 | 3'UTR | -100 | 94 | 20 |
| *MAP7* | miR529a-3p | 2137 | CDS | -100 | 94 | 20 |
| *MLL* | miR529a-3p | 11418 | CDS | -100 | 94 | 20 |
| *PDE4B* | miR529a-3p | 2270 | CDS | -100 | 94 | 20 |
| *ENAH* | miR531b-5p | 106 | 5'UTR | -115 | 95 | 20 |
| *TIFA* | miR815a,b,c-3p | 186 | 5'UTR | -106 | 94 | 21 |

The miRNAs of the large miR395-3p family had *EDN1* and *CD22* target genes, which are involved in the development of diabetes and in the control of immunity, respectively. If these miRNAs get in food in large quantities, then the probability of their impact on human health is high.

miR396a,b-3p can affect the expression of the *CMKLR1* gene, which is involved in cardiovascular disease, and for miR396a,b-5p, the *SCAMP5* and *SYVN1* genes are targeted, the expression of which changes with autism and colon cancer, respectively. miRNAs of the miR444-3p family have binding sites in the mRNA of six genes (Table S4). Their target genes *RHBDF1, RUNDC1, CEP250, C19orf57, KAZN*, and *NRG1* are involved in oncogenesis and other diseases.

Therefore, ingestion of these miRNAs with food in humans can significantly affect metabolic processes. miR529a-3p had binding sites in the mRNA of six genes that are involved in the regulation of several physiological processes (Table S5). If a person in the process of evolution consumed this miRNA as a necessary regulator of the expression of its target genes, then for this miRNA there must be target genes.

Fig. 3 shows the interaction patterns of the nucleotide sequences of some representatives of the miRNA families with the mRNA of their target genes. These data indicate a good predictive power for identifying miRNA associations and target genes. The visibility of the interaction of miRNA and mRNA nucleotides in combination with the quantitative characteristics of binding miRNA and mRNA allows us to consider these associations stable and real.

**Figure 3.**
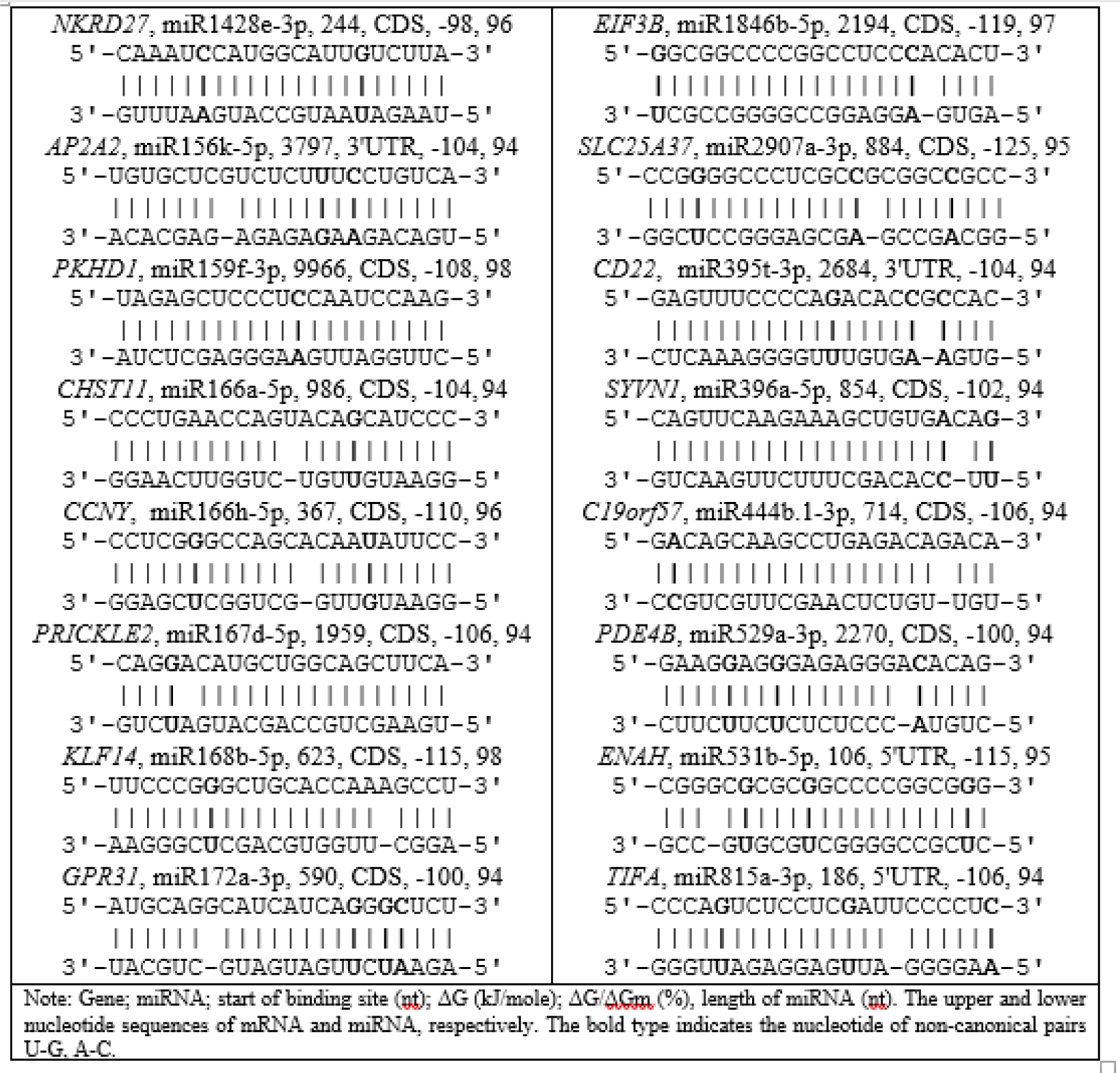
Schemes of the interaction of nucleotide sequences of osa-miRNA families with mRNA human genes.

Note that the nucleotide sequences miR156, miR166, miR395, miR396, and miR444 did not have homologous miRNAs among 2565 human miRNAs from the miRBase base. Therefore, these miRNAs do not directly have common binding sites for human miRNAs and can independently regulate the expression of their target genes.

To confirm the conservatism of the interaction of pl-miR with human target genes, we plotted the web logo for mRNA sections containing pl-miR binding sites (Fig. 2). The graphs show the high conservatism of these binding sites compared to flanking nucleotide sequences. The miR156a-j-5 family, consisting of 10 miRNAs, was associated with the mRNA of seven genes (Table S4). The miR156a-j-5 binding sites in each of the genes differed in the number of hydrogen bonds (Fig. 3) and the value of the free interaction energy. Similar results were obtained for other miRNA families: miR164e-5p, miR168b-5p, miR396c-3p, miR444a-3p.1,d.1-3p, miR529a-3p, miR815a,b,c-3p, miR1846a,b,c-5p, miR1858a,b-5p, miR2118l-3p, miR2275d-3p, miR2907a,b,d-3p, and miR395a-y-3p (Fig. 2). The conservatism of such bonds between miRNAs and their target genes was established by us for many associations of miRNAs and their target genes in animals and plants (Atambayeva et al., 2017; Bari, Orazova & Ivashchenko, 2013; Bari et al., 2014; Yurikova et al., 2019). These bonds have persisted over tens of millions of years of evolution and indicate the early emergence of the process of regulation by miRNA molecules of target gene expression in animals and plants.

## Discussion

Based on the results obtained in this work, the fact of the interaction of pl-miRs with mRNA of human genes is beyond question. It is necessary to establish the possibilities for these miRNAs to enter the human and animal organisms. Several studies have shown that miRNAs in various parts of plants are present in exosomes 30-400 nm in size and are distributed in the body as part of these nanoparticles (Bang & Thum, 2012; Denzer et al., 2000; Xiao et al., 2018). Such compaction of miRNAs in exosomes contributes to their conservation and facilitates the entry of miRNAs into animals through the digestive tract (Redis et al., 2012; Théry, Zitvogel & Amigorena, 2002; Valadi et al., 2007). Further exosomes together with endogenous exosomes with blood move to many tissues and organs. According to the physicochemical properties, plant miRNAs do not differ from animal miRNAs, which makes them competitive when interacting with mRNA target genes. There are no known limitations for the above-described process of ingestion of plant-miRNAs into humans and animals.

Basically, not all pl-miRs will have human target genes, but the most common and vital pl-miRs present in plants can have target genes in animals and humans for a long time eating them. Such pl-miRs usually participate in maintaining the basic physiological functions of plants (productivity, resistance to biotic and abiotic stresses, growth and development). For example, developed rice lines overexpressing *MIR529a* have been shown to have increased resistance to oxidative stress (Chen & Li, 2018; Cimini et al., 2019). The participation of osa-miR159f, osa-miR1871, osa-miR398b, osa-miR408-3p, osa-miR2878-5p, osa-miR528-5p and osa-miR397a in the regulation of a number of physiological processes of rice has been established (Balyan et al., 2017). The expression of miRNA of *Setaria italica* (sit) changed many times: sit-miR1432-3p, sit-miR156a-5p, sit-miR156b-5p, sit-miR164a-5p, sit-miR167b-5p, sit-miR171c-3p, sit-miR2118-3p, sit-miR390-5p, sit-miR394-5p, sit-miR395-3p, sit-miR408-3p, sit-miR529a-3p, sit-miR529b-3p, and sit-miR827, sit-miR159b-3p, sit-miR319c-5p, sit-miR528-5p and sit-miR535-5p under various stresses (Wang et al., 2016). In a broader evolutionary context, miRNAs of *Morus notabilis* were compared to those of seven other plants, including five dicotyledons, *Arabidopsis thaliana*, *Glycine max*, *Malus domestica*, *Populus trichocarpa*, *Ricinus communis*, and two monocotyledons, *Oryza sativa* and *Zea mays*. Of the 31 *Morus notabilis* miRNA families, 24 were conserved in the seven plant species. These miRNAs were classified into well-conserved miRNA families. Prominent among them were mulberry miR160b, miR164a, miR167a, miR169a, miR390, and miR396b, which completely matched their counterparties in the seven other plant species, suggesting that those miRNAs were extremely conserved, and might play critical physiological roles in both dicotyledons and monocotyledons. However, seven miRNA families, miR482, miR529, miR858, miR4376, miR4414, miR4995, and miR5523, were found in only one or two plant species. The present data indicated that the conserved miRNA families (miR156, miR166, miR167, miR168, and miR535) miR159, miR160, miR164, miR169, miR171, miR172, miR390, miR396, miR397, miR529 and miR4376, miR162, miR393, miR395, miR398, miR399, miR408 and miR4414 miR319, miR482, miR827, miR828, miR858, miR2111, miR4995 and miR5523 were expressed across a vast range exceeded in all three tissues (Jia et al., 2014). In African rice *Oryza glaberrima (*ogl), some miRNAs such as ogl-miR156l, ogl-miR166c, ogl-miR166k, ogl-miR168a, ogl-miR167i, ogl-miR171f, ogl-miR1846d of the control library and ogl-miR408, ogl-miR528, ogl-miR156, ogl-miR390, and ogl-miR396c of the treated library had higher reads than their complementary strand. This is because miRNA-3p and miRNA-5p may function simultaneously to regulate gene expression (Mondal et al., 2018). It must be understood that the expression of the target gene under the influence of miRNAs can increase with decreasing miRNA concentration below the average physiological level or decrease with increasing miRNA concentration. The data on changes in miRNA concentration in different plants show that the amount of miRNA consumed with food depends on the stage of plant ontogenesis, growing conditions, plant organs, food processing, etc. (Liu et al., 2017). The concentration of pl-miRs after various processing of raw products decreases, but the remaining miR enter the body (Luo et al., 2017; Zhang et al., 2012; Zhou et al., 2015).

## Conclusions

As a result of our study, for the first time, among 17508 human genes, 942 target genes for 277 osa-miRNAs were established. The identified target genes account for 5.4% of the total number of studied human genes. miRNA binding sites were found in the CDS, 5’UTR and 3’UTR. The largest number of genes were targeted by osa-miR2102-5p, osa-miR5075-3p, osa-miR2097-5p, and osa-miR2919, which can bind to the mRNA of 38, 36, 23, and 19 genes, respectively. Since osa-miRNAs ingested through plant food have many target genes, they should be controlled in the human body. Most osa-miRNA target genes are involved in the development of diseases, which makes it easier to clarify the role of miRNAs in these processes. Many osa-miRNA target genes contribute to the development of breast cancer, and other cancer types. The other target genes are involved in cardiovascular and neurodegenerative diseases. Some osa-miRNAs can be effective regulators of human gene expression. The effect of miRNAs can be both positive, contributing to the cure of diseases, and negative, causing a wide range of diseases.

## Materials & Methods

The nucleotide sequences of the mRNAs of 17508 targeted genes were downloaded from NCBI GenBank (http://www.ncbi.nlm.nih.gov). The nucleotide sequences of the miRNAs were taken from miRBase v.22 (http://www.mirbase.org/). The miRNA binding sites in the mRNAs of several genes were predicted using the MirTarget program (Ivashchenko et al., 2016). This program defines the following features of miRNA binding to mRNA: a) the start of the initiation of the miRNA binding to the mRNAs from the first nucleotide of the mRNA’s; b) the localization of the miRNA binding sites in the 5’-untranslated region (5’UTR), coding domain sequence (CDS) and 3’-untranslated region (3’UTR) of the mRNAs; c) the free energy of the interaction between miRNA and the mRNA (∆G, kJ/mole); and d) the schemes of nucleotide interactions between miRNAs and mRNAs. The ratio ΔG/ΔGm (%) is determined for each site (ΔGm equals the free energy of the miRNA binding with its fully complementary nucleotide sequence). The MirTarget program finds hydrogen bonds between adenine (A) and uracil (U), guanine (G) and cytosine (C), and G and U, A and C. The distances between the bound A and C (1.04 nm) and G and U (1.02 nm) are similar to those between bound G and C and A and U, which are equal to 1.03 nm (Garg & Heinemann, 2018; Kool, 2001; Leontis, Stombaugh & Westhof, 2002). The numbers of hydrogen bonds in the G-C, A-U, G-U and A-C interactions were 3, 2, 1 and 1, respectively. By comparison, MirTarget differs from other programs in terms of finding the binding sites of miRNA on the mRNAs of plant genes (Dai, Zhuang & Zhao, 2011) in that 1) it takes into account the interaction of the miRNA with mRNA over the entire miRNA sequence; 2) takes into account noncanonical pairs G-U and A-C; and 3) calculates the free energy of the interaction of the miRNA with mRNA, and when two or more miRNAs are bound with one mRNA or, if the binding sites of two different miRNAs coincide in part, the preferred miRNA binding site is considered to be the one for which the free binding energy, ∆G, is greater. The MirTarget program does not work directly with the miRBase and NCBI databases. The search for target genes from 17,508 human genes in a special format from NCBI for the known miRNAs from miRBase will be available on request at.

## Suplemental Data

**Supplemental table 1.** osa-miRNA list of 1-4 human target genes.

**Supplemental table 2.** osa-miRNA list of 5 or more human target genes.

**Supplemental table 3.** osa-miRNA of target human genes involved in biological processes.

**Supplemental table 4.** list of osa-miRNA families of human target genes.

**Supplemental table 5.** osa-miRNA families of target human genes involved in biological processes.

## AUTHOR CONTRIBUTIONS

A.I., A.R performed the research and analyzed the data. A.P., D.A contributed analytic tools and methods. A.I., A.R., A.P., D.A wrote the article.

